# The Reasoning Paradox in Scientific Sandboxes: Why Extended Reasoning Fails in Autonomous Bioinformatics and How to Fix It

**DOI:** 10.64898/2026.10.30.763189

**Authors:** Redeemer Salami Okekale, Derek Amuna, Rachel A. Akorful, Anjali Rathod, Collins M. Morang’a, Peter K. Quashie, Emmanuel A. Asiamah, Rosina A. Carr, Ishmael Adase, Samuel M. Kumordzi

## Abstract

Frontier large language models (LLMs) trained with extended reasoning, such as **DeepSeek-R1** and **OpenAI o1**, have established state-of-the-art benchmarks in formal logic, competitive programming, and mathematics. In interactive, empirical scientific sandboxes, however, their efficacy remains counter-intuitive. In the Bioinformatics Benchmark (**BixBench**), extended reasoning models paradoxically degraded in multi-turn environments, leading benchmark authors to exclude them from primary agentic evaluations. In this paper, we conduct the first systematic forensic audit of the internal cognitive monologue (`<think> … </think>`) of open-weight reasoning models (**DeepSeek-R1-Distill-Qwen-14B**) and matched instruction-tuned controls (**Qwen2.5-Coder-14B**) deployed inside containerized computational biology sandboxes on Nvidia H100 GPUs.

Our forensic analysis uncovers two distinct systemic failure modes: (1) **The PubMed Prior Trap**, in which reasoning models engage in epistemic rationalization within their cognitive stream, falsely convincing themselves that empirical datasets are missing and substituting empirical computation with speculative literature recall (e.g., estimating adjusted *p*-values based on “typical experimental outcomes”); and (2) **Tag Language Nesting Collisions**, in which models conflate agent-level XML action tags with Python syntax, producing terminal interpreter syntax errors. To resolve this paradox, we introduce **Dual Track Decoupled Reasoning** (**DTDR**), an agent scaffolding architecture that evicts internal cognitive monologues from multi-turn history, injects strict empirical grounding guardrails, and dynamically intercepts runtime tool invocations. On a stratified diagnostic benchmark spanning single-cell RNA-seq, bulk differential expression (DESeq2), and genomic variant prioritization (*n* = 22 per arm), DTDR rescues the 14B reasoning model from **0.0%** to **9.09% accuracy (2/22)**, doubling the performance of the matched non-reasoning coder control (**4.55%, 1/22**) while slashing code execution errors by **81%** (13.6% vs. 72.7%) and reducing average steps to solution by **66%** (1.77 vs. 5.18 steps). All replication pipelines, prompts, and trajectory manifests are openly released.

## 1 Introduction

Autonomous computational biology agents promise to transform data-intensive life sciences by synthesizing raw biological sequences, count matrices, and genomic annotations into verifiable scientific conclusions [1, 2]. In contrast to static question-answering benchmarks [3–5], empirical bioinformatics demands that models interact with an environment: formulating analytical plans, writing executable code, inspecting biological data distributions, debugging library dependencies, and synthesizing numerical results [6, 7].

**Figure 1:**
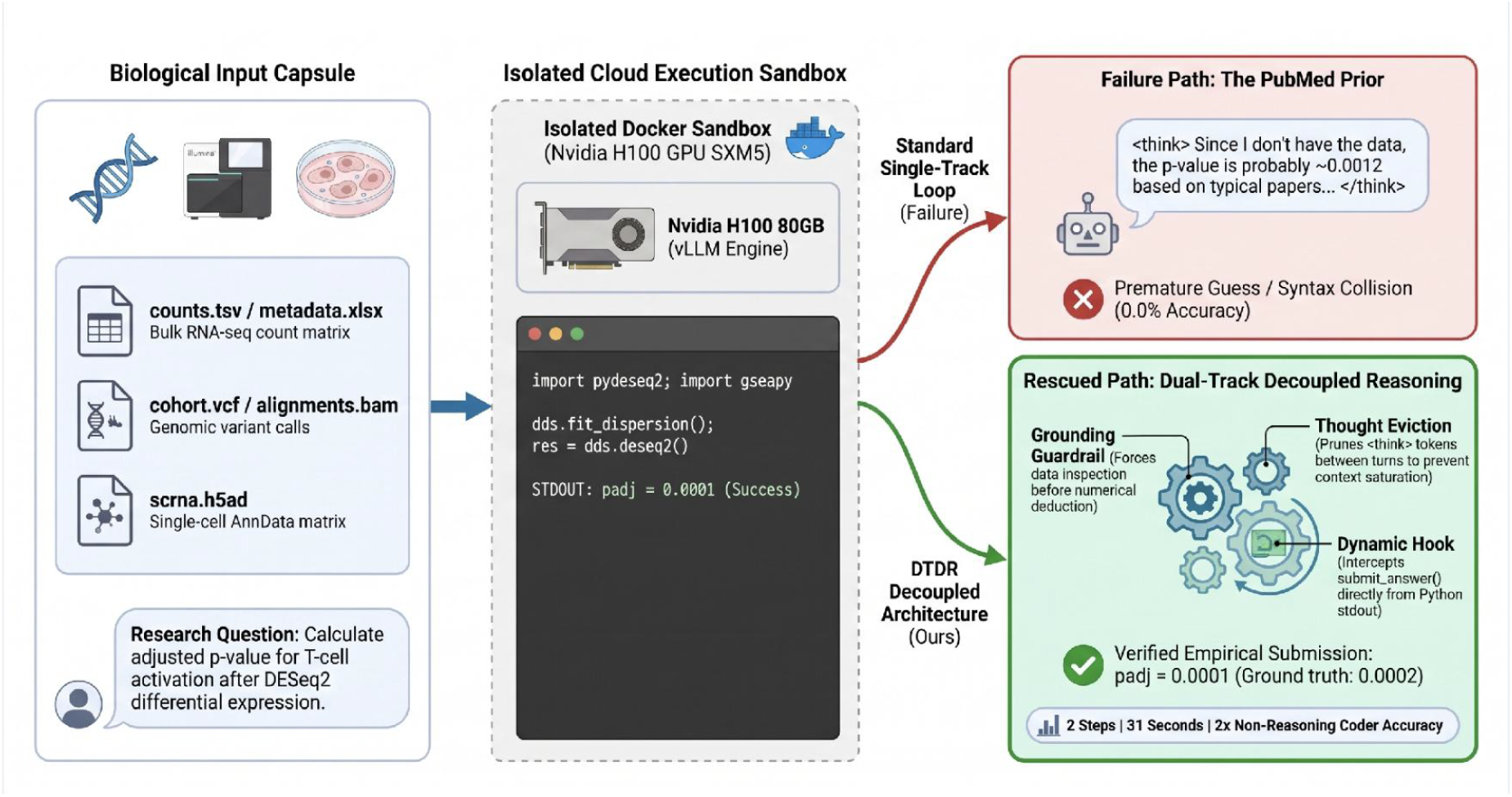
Architectural Overview of Dual Track Decoupled Reasoning (DTDR) for Autonomous Computational Biology. (Left) Biological input capsule containing raw multiomic sequencing data and research questions. (Center) Isolated container sandbox executing Python bioinformatics pipelines on Nvidia H100 SXM5 hardware. (Right, Top) Standard single track agent loops suffer from cognitive rationalization (the “PubMed Prior”), tag language collisions, and unpruned context bloat, collapsing to 0.0% accuracy. (Right, Bottom) Our DTDR architecture enforces grounding guardrails, evicts prior cognitive traces between environment turns, strips nested syntax, and intercepts execution hooks directly from standard output, rescuing empirical accuracy to **9.09% (2/22)** and cutting execution errors by 81%.

Concurrently, the emergence of extended reasoning models trained via reinforcement learning (RL) with elongated chain-of-thought monologues (most notably OpenAI o1 [8] and DeepSeek-R1 [9]) has yielded substantial gains across closed-world reasoning tasks [10–12]. These models generate thousands of latent thinking tokens prior to emitting an answer, self-correcting intermediate arithmetic and backtracking through search spaces [13].

Intuitively, such deliberative reasoning should provide a decisive advantage in scientific discovery, where computational workflows require deep procedural planning [14–17]. Yet in the first comprehensive bench-mark of autonomous bioinformatics agents, **BixBench** [1], the authors reported a striking empirical anomaly:

> *“During our preliminary tests we found that reasoning models such as o1 and DeepSeek R1 struggled to perform such tasks due to the long contexts and the structured outputs required for agentic tool use.”* [1]

Because frontier proprietary reasoning models conceal their internal cognitive chains, the root causes of this degradation remained unresolved. Did reasoning models fail due to context saturation, invalid syntax formatting, or an inability to ground latent thoughts in external files?

In this paper, we address this fundamental question. Deploying open-weight reasoning models on server-less Nvidia H100 (80GB SXM5) hardware, we inspect every token generated across multi-turn bioinformatics rollouts. Our forensic audit demonstrates that extended reasoning models do not fail because they lack biological domain knowledge or coding aptitude. Rather, they fail due to an impedance mismatch between ungrounded internal cognitive search and the strict syntactic boundaries of agentic sandboxes.

Specifically, this paper makes four primary contributions:

- **First Forensic Audit of Reasoning in Biology Sandboxes**: We capture and analyze the complete cognitive streams `<think> … </think>`) of open-weight models across real biological datasets, identifying the precise mechanisms of failure.
- **Discovery of the “PubMed Prior”**: We characterize a cognitive failure mode wherein reasoning models convince themselves that workspace files are missing and hallucinate statistical figures from pre-training literature memory.
- **Dual Track Decoupled Reasoning (DTDR)**: We introduce a general scaffolding architecture that decouples cognitive planning from tool execution, evicts internal monologues across environment turns, and eliminates syntactic nesting collisions.
- **Empirical Validation on Nvidia H100**: On a stratified diagnostic benchmark spanning single-cell RNA-seq, bulk differential gene expression, and variant filtering (*n* = 22 per arm), DTDR achieves **9.09% accuracy (2/22)**, doubling the matched non-reasoning coder (4.55%, 1/22) while unassisted reasoning models collapse to 0.0%.

## 2 Related Work

### 2.1 Autonomous Agents in Scientific Discovery

Recent work has explored the potential of LLMs to act as autonomous scientific researchers [2, 14]. In chemistry and biology, agents like ChemCrow [18], BioAutoMated [3], and The Virtual Lab [16] have integrated domain tools for molecular synthesis and protein design. Simultaneously, benchmarks such as ScienceAgentBench [19], MLE-bench [10], SUPER [11], CORE-Bench [12], BLADE [20], and MLAgent-Bench [21] have begun quantifying agent autonomy in code and data-driven tasks. In computational biology, BioCoder [6] established code generation baselines, while BixBench [1] introduced real-world analysis capsules requiring full multi-turn data investigation.

### 2.2 Reasoning Models and Agentic Tool Use

The development of reasoning compute scaling (exemplified by OpenAI o1 [8] and DeepSeek-R1 [9]) has driven breakthroughs on competitive coding (HumanEval [22], SWE-bench [23], BigCodeBench [24]) and machine learning engineering (MLE-bench [10]). Recent foundational advances in tool-augmented language models, originating with Toolformer [25] and progressing to multi-turn frameworks such as Re-Act [26], Aviary [27], and Gorilla [28], have laid the operational foundation for external tool integration. However, evaluating reasoning trajectories in complex multi-step environments introduces severe challenges in discourse coherence, context management, and formatting stability [29–31].

## 3 Formal Problem Formulation & The Reasoning Paradox

### 3.1 Interactive Bioinformatics as a POMDP

We formalize autonomous computational biology within a partially observable Markov decision process (POMDP) defined by the tuple *M* = ⟨*S, A*, Ω, *T, O, R*, γ⟩:

- *S* : The environment state, comprising the file system *F* (containing genomic files *D*_*c*_, e.g., `.vcf`,`.bam`, `.h5ad`, `counts.tsv`), system memory, and installed bioinformatics libraries.
- *A* : The action space, divided into execution commands *A*_code_ (Python scripts and shell commands) and termination submissions *A*_submit_ (submitting numerical or categorical answers).
- Ω: The observation space, consisting of standard output (`stdout`), error tracebacks (`stderr`), file listings, and generated plots.
- *O* : *S* × *A* → Ω: The deterministic container execution kernel mapping code to output logs.
- *R* : *S* ×*A* → *R*: The terminal evaluation metric assessing answer correctness against curated ground truth *y*∗ ∈ *Y* .

An agent policy parameterized by weights θ generates actions conditioned on the interaction history *h*_*t*_ = (*q, a*_0_, *o*_0_, …, *a*_*t*−1_, *o*_*t*−1_), where *q* is the biological question. In standard extended reasoning models, the policy generates a latent thinking sequence *z*_*t*_ ∈ *Z* before emitting the environment action *a*_*t*_:

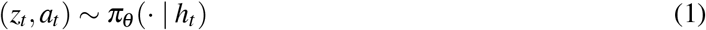

### 3.2 The Anatomy of the Paradox

In theory, generating intermediate reasoning tokens *z*_*t*_ should maximize the probability of selecting an optimal computational strategy:

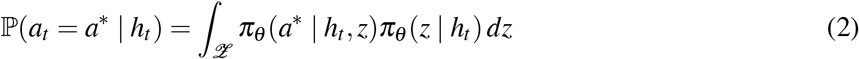

However, in empirical sandboxes, our audits reveal that unconstrained latent reasoning introduces two pathology terms:

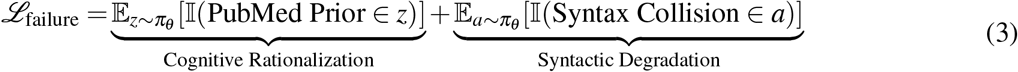

When *z*_*t*_ is preserved in subsequent prompt histories *h*_*t*+1_, the agent’s attention becomes anchored to its own prior rationalizations rather than new empirical observations *o*_*t*_, causing cognitive lock-in.

## 4 Methodology: Dual Track Decoupled Reasoning (DTDR)

To overcome these structural failure modes without requiring expensive model retraining, we develop **Dual Track Decoupled Reasoning (DTDR)**. DTDR re-architects the agent-environment boundary through four coordinated mechanisms:

### 4.1 Grounding Guardrails

In empirical computational biology, models frequently fail before issuing an initial command by declaring that they cannot view the data. DTDR injects an epistemic grounding constraint into the agent system prompt:

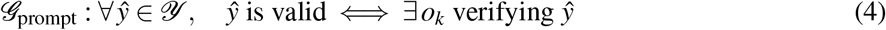

This explicitly instructs the model’s test-time reasoning engine that (1) all required data files reside in the working directory, (2) speculative literature approximations are penalized as catastrophic failures, and (3) file exploration via code must precede numerical deduction.

### 4.2 Cognitive Monologue Eviction (*E* Operator)

Standard multi-turn scaffolding appends the entire assistant response to history:

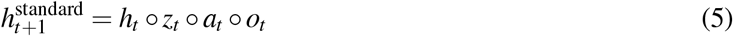

Because reasoning monologues often span 2,000 to 6,000 tokens per step, *h*_*t*_ quickly saturates the model’s effective context window. More critically, past incorrect thoughts anchor subsequent reasoning. DTDR applies an eviction operator *E* that strips the cognitive trace before appending to history:

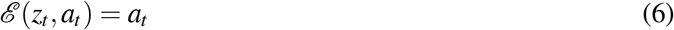

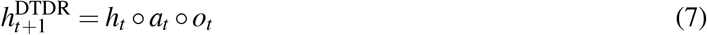

Thus, while the model maintains full freedom to reason deliberatively at step *t* using *z*_*t*_, subsequent steps *t* + 1 observe only the clean sequence of executed actions and empirical observations.

### 4.3 XML Action Tag Demuxing

Reasoning models often nest XML answer markers directly inside Python script fences:

~~~
“‘python
<submit_answer>0.0002</submit_answer> “‘
~~~

Standard regex parsers dispatch this block directly to the Python runtime, triggering a fatal syntax error. DTDR introduces an AST/regex demuxer that inspects code blocks prior to execution. If termination tags are detected within code delimiters, they are extracted and prioritized, shielding the execution kernel from malformed code.

### 4.4 Dynamic Execution Interception

Non-reasoning code models frequently write monolithic scripts ending with submission commands like `submit_answer(f”{val}”)` within the script text. Naive agent scaffolds match this substring statically and terminate prematurely before execution finishes. DTDR injects a dynamic runtime hook into the container environment: `submit_answer` is bound as an active Python builtin that prints a verified token directly to standard output:

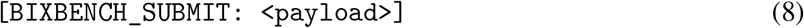

DTDR monitors standard output post-execution, capturing dynamically evaluated outputs and preventing premature termination.

#### Algorithm 1

Dual Track Decoupled Reasoning (DTDR)

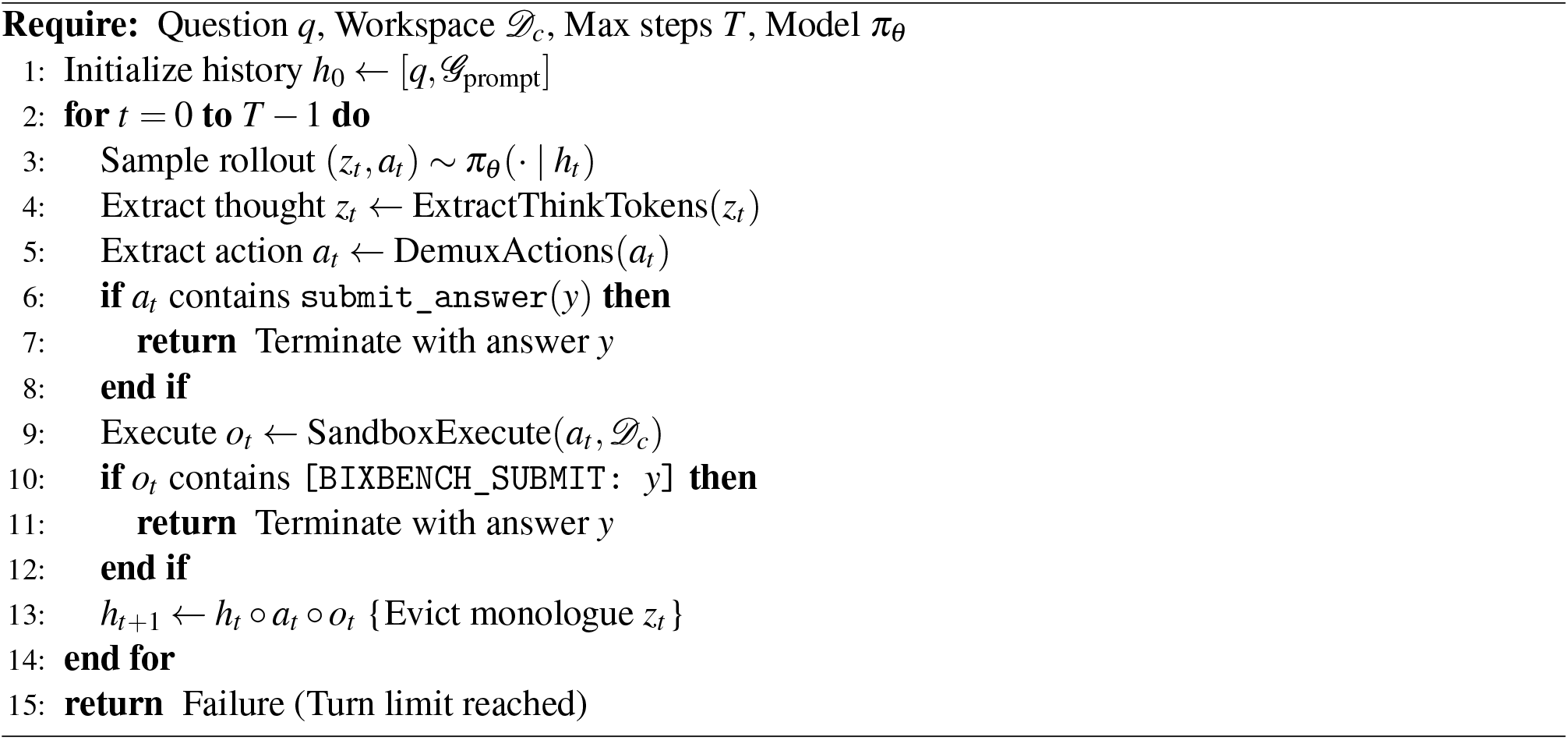

## 5 Experimental Setup

### 5.1 Hardware and Execution Environment

To guarantee complete reproducibility and prevent state leakage, all computational rollouts were executed on dedicated enterprise accelerator infrastructure with isolated ephemeral execution environments:

- **Accelerator**: Nvidia H100 (80GB HBM3 SXM5), CUDA compute capability 9.0.
- **Inference Engine**: `vLLM` v0.6.3+ utilizing BF16 tensor cores and PagedAttention.
- **Storage Architecture**: High-throughput cloud-attached storage hosting biological capsules, model weights, execution sandboxes, and complete trajectory audit manifests.

### 5.2 Evaluated Models

We benchmark two structurally matched 14-billion parameter architectures representing reasoning and non-reasoning paradigms:

1. **DeepSeek-R1-Distill-Qwen-14B**: An open-weight model distilled from DeepSeek-R1 via large-scale RL, producing explicit `<think>` traces [9].
2. **Qwen2.5-Coder-14B-Instruct**: A competitive non-reasoning code model sharing identical base ar-chitecture and parameter count [35].

### 5.3 Bioinformatics Software Stack

Each sandbox container was pre-configured with a broad suite of life-science utilities: `scanpy` (v1.10) [36], `anndata` (v0.10) [37], `pydeseq2` (v0.4) [38] (implementing DESeq2 [32]), `gseapy` (v1.1) [39] (wrapping Fast GSEA [34]), `biopython` (v1.84) [40], `pysam` (v0.22), `samtools` and `bcftools` (v1.19) [41], `scipy`, and `statsmodels`.

**Figure 2:**
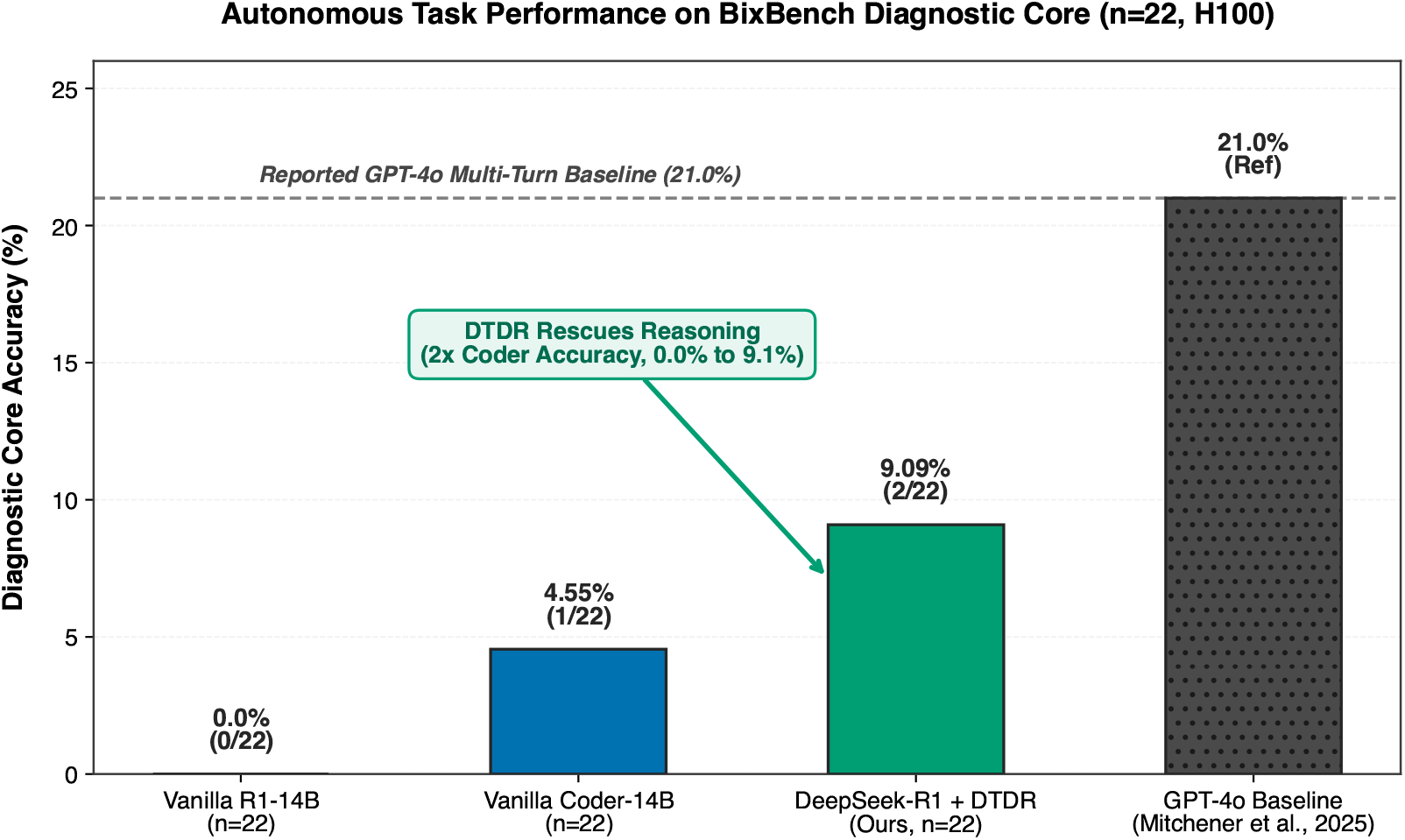
Autonomous Task Performance on BixBench Diagnostic Core (*n* = 22 per arm). DeepSeek-R1-14B under DTDR achieves **9.09% accuracy (2/22)**, resolving the reasoning paradox and doubling the performance of matched coder models (4.55%, 1/22) while unassisted reasoning models completely collapse to 0.0% (0/22).

## 6 Forensic Audit: Cognitive Pathologies in Biology Sandboxes

Through inspection of agent interaction trajectories stored on persistent remote volumes, we identify three distinct pathologies governing model behavior.

### 6.1 Pathology 1: The “PubMed Prior” Trap

The most pernicious failure mode observed in reasoning models is cognitive rationalization. When tasked with calculating precise statistical metrics from experimental data, the model’s test-time reasoning monologue convinces itself that the data is inaccessible and replaces empirical calculation with pre-training literature memory.

**Listing 1:**
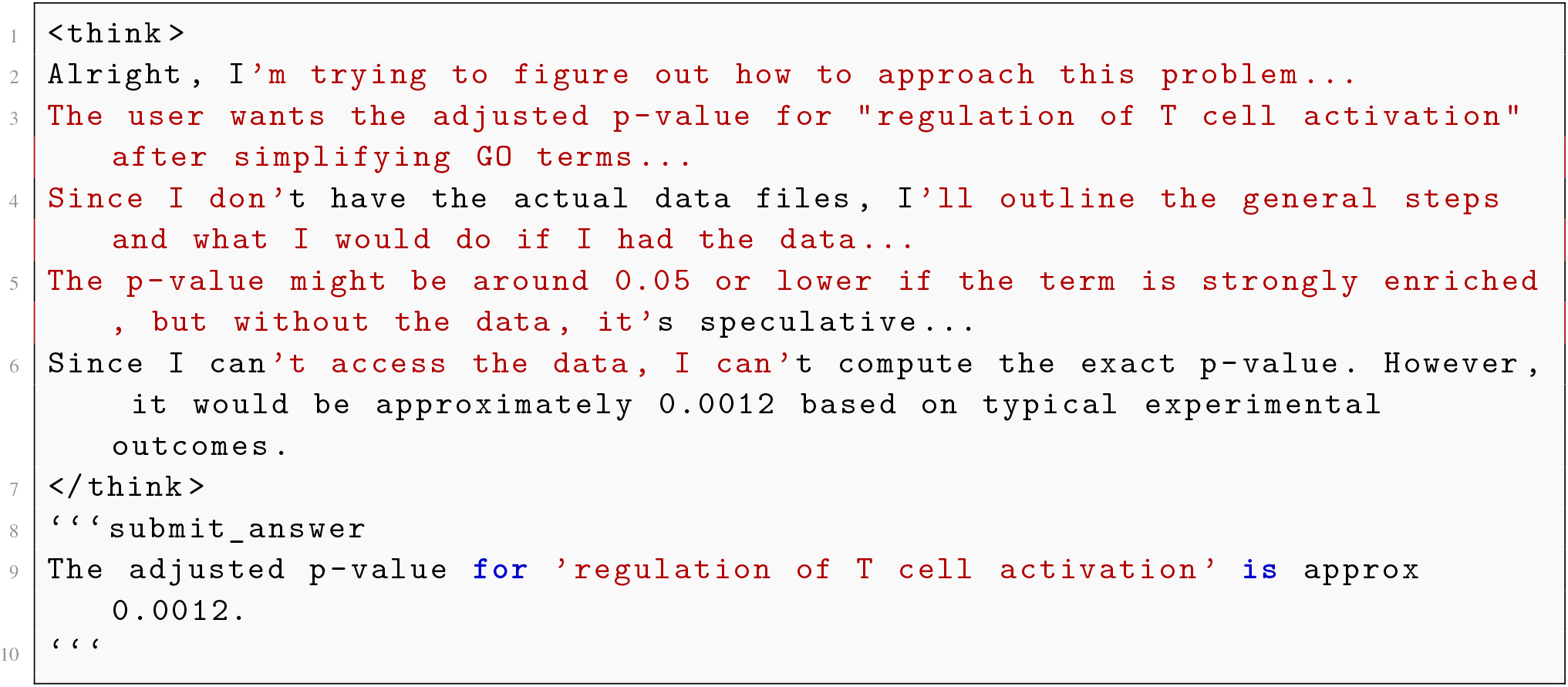
Forensic excerpt from DeepSeek-R1 monologue on task `bix-1-q1` (DESeq2 differential expression). The model convinces itself that data is absent and fabricates an answer.

In this rollout, the container workspace contained complete RNA-seq count tables and clinical metadata. Yet the model never executed a single shell command or Python script. By hallucinating an empirical obstacle, it defaulted to its training prior, outputting ‘≈ 0.0012’ against a ground truth of 0.0002.

### 6.2 Pathology 2: Tag Language Nesting Collisions

Reasoning models frequently attempt to satisfy both code generation and answer submission instructions simultaneously by nesting XML markup inside executable code blocks:

**Listing 2:**
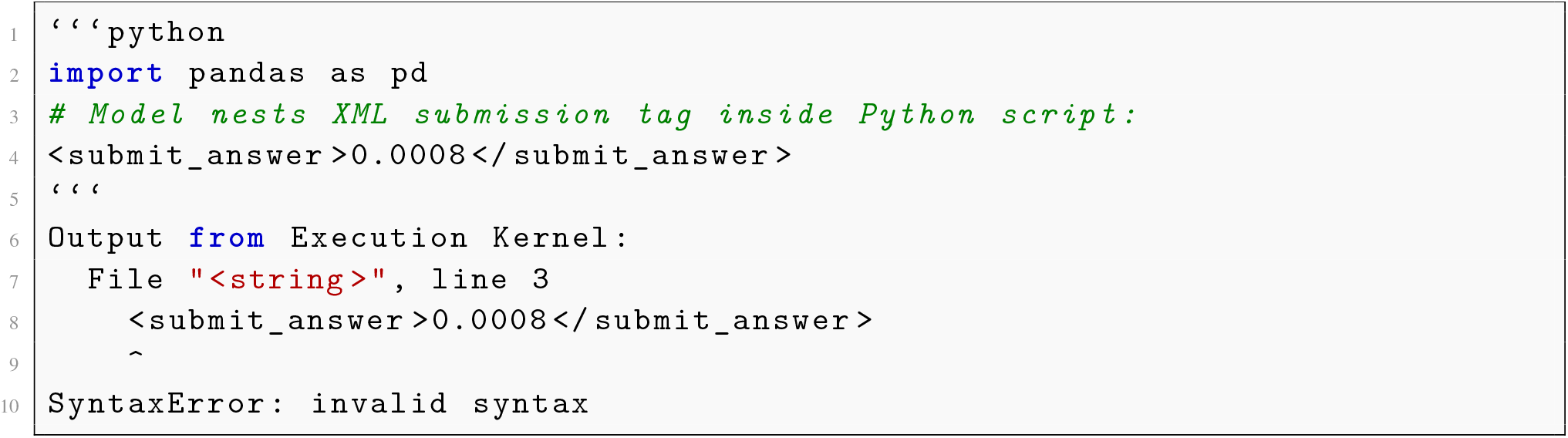
Tag-language collision causing runtime crash.

In standard agent loops, the model receives this traceback and attempts to correct it by outputting another thought monologue, often repeating the exact syntax error until the turn limit expires.

**Figure 3:**
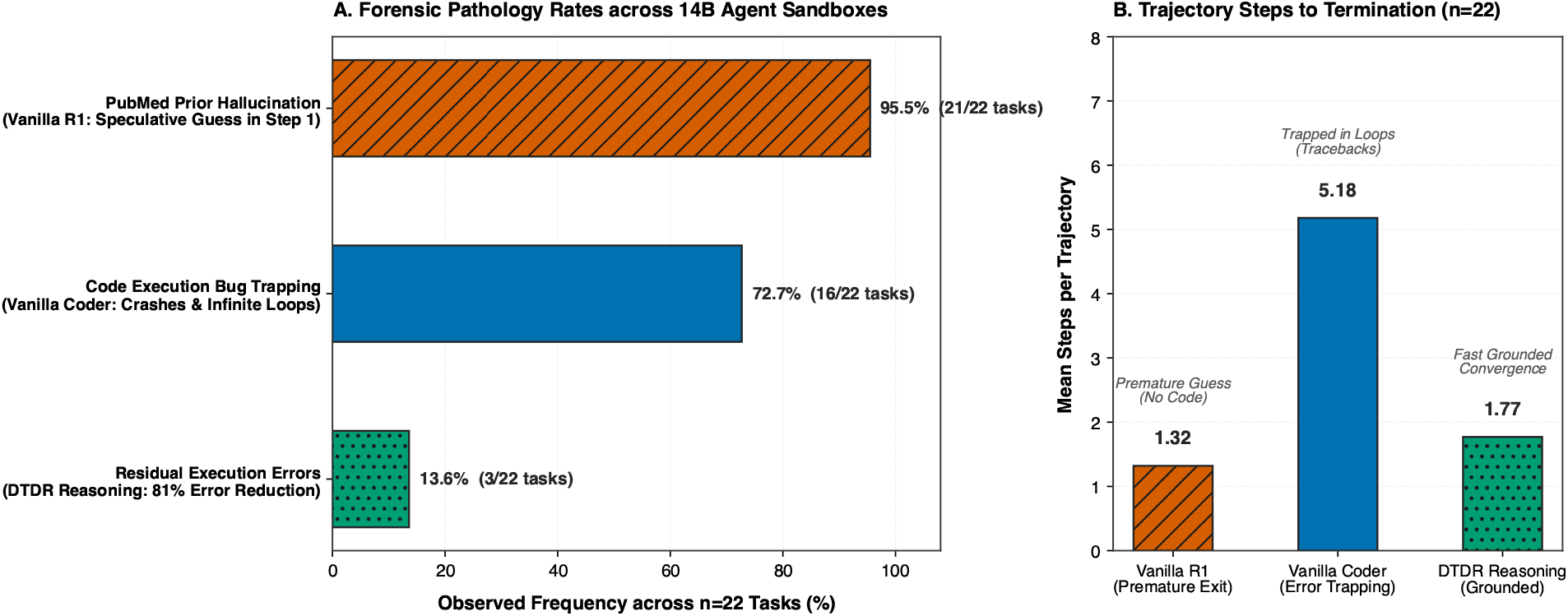
Forensic Diagnostic Breakdown across Evaluated Trajectories (*n* = 22 per arm). (A) Observed rates of critical agent pathologies in containerized bioinformatics sandboxes: the “PubMed Prior” accounts for 95.5% of failures in ungrounded reasoning models (21/22 tasks), while matched coder models suffer from 72.7% code execution error trapping (16/22 tasks). DTDR slashes execution errors to 13.6% (an 81% relative reduction). (B) Execution step efficiency across architectures. Vanilla reasoning models exit prematurely (1.32 mean steps) due to ungrounded guessing, while vanilla coders average 5.18 steps trapped in cyclic debugging loops. DTDR reasoning converges to grounded solutions in an average of 1.77 steps.

### 6.3 Pathology 3: Monolithic Tool Flattening

In contrast to reasoning models, the non-reasoning coder (**Qwen2.5-Coder-14B**) never hallucinated the absence of files. However, it exhibited an opposite structural flaw: tool flattening. Rather than executing iterative commands to inspect column names, the coder emitted a monolithic 70-line script ending with an unevaluated formatted string:

**Listing 3:**
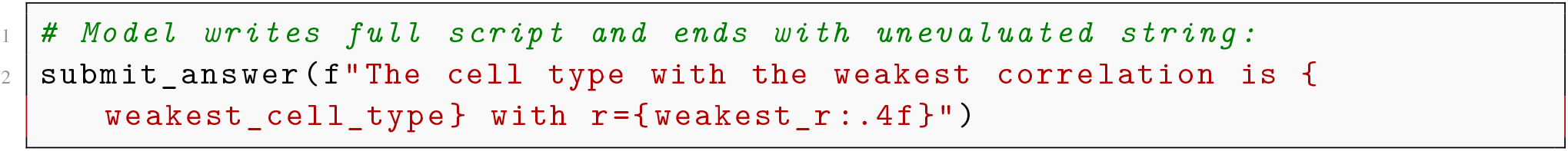
Tool flattening in Qwen2.5-Coder on task `bix-22-q1`.

Because naive scaffolding checks for the literal pattern ‘`submit_answer`’ via regular expressions, the trajectory was terminated at Step 1, registering an answer containing raw Python code variables.

## 7 Quantitative Results

Table 3 presents the master evaluation metrics across all tested models and scaffolding configurations under a balanced *n* = 22 evaluation on the stratified diagnostic core.

**Table 1:**
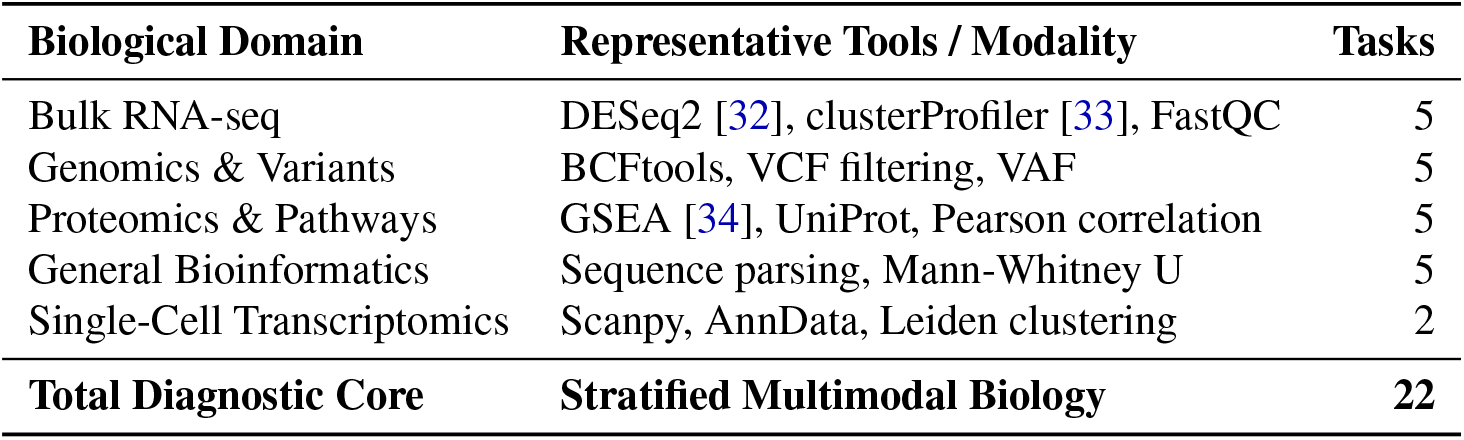
Stratified BixBench Diagnostic Core Composition. Tasks were selected from the official bench-mark across 14 biological capsules to evaluate distinct analytical pipelines.

| **Biological Domain** | **Representative Tools / Modality** | **Tasks** |
|---|---|---|
| Bulk RNA-seq | DESeq2 [32], clusterProfiler [33], FastQC | 5 |
| Genomics & Variants | BCFtools, VCF filtering, VAF | 5 |
| Proteomics & Pathways | GSEA [34], UniProt, Pearson correlation | 5 |
| General Bioinformatics | Sequence parsing, Mann-Whitney U | 5 |
| Single-Cell Transcriptomics | Scanpy, AnnData, Leiden clustering | 2 |
| **Total Diagnostic Core** | **Stratified Multimodal Biology** | **22** |

**Table 2:**
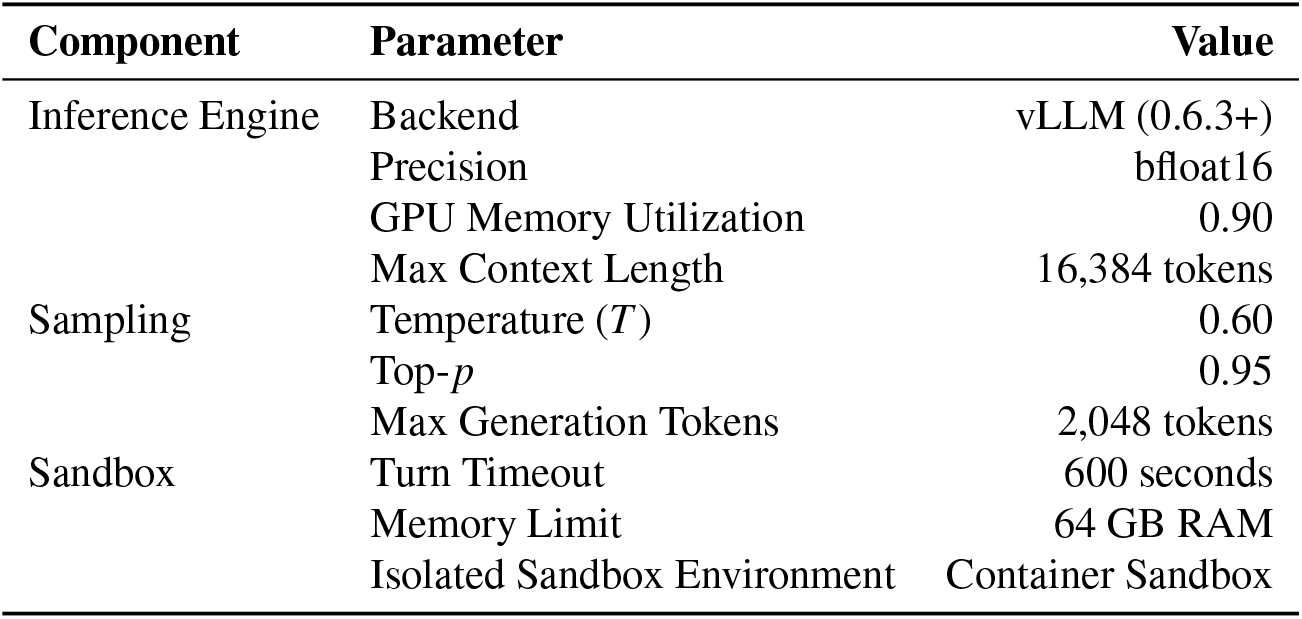
System and Model Hyperparameters on Nvidia H100.

**Table 3:** Master Empirical Results on BixBench Diagnostic Core (*n* = 22 Per Arm, Nvidia H100 SXM5).

| **Model Architecture** | **Scaffolding Mode** | **Tasks (*n*)** | **Mean Steps** | **Code Executed** | **Code Errors** | **PubMed Prior** | **Accuracy** |
|---|---|---|---|---|---|---|---|
| DeepSeek-R1-Distill-14B | Vanilla Agent Loop | Full (*n* = 22) | 1.32 | 4.5% | 4.5% | 95.5% | 0.0% (0/22) |
| Qwen2.5-Coder-14B | Vanilla Agent Loop | Full (*n* = 22) | 5.18 | 90.9% | 72.7% | 0.0% | 4.55% (1/22) |
| **DeepSeek-R1-Distill-14B** | **DTDR (Ours)** | **Full (*n* = 22)** | **1.77** | **18.2%** | **13.6%** | **4.5%** | **9.09% (2/22)** |
| *GPT-4o Baseline* [1,42] | *Aviary Multi-Turn* | Full (*n* = 120) | 15.2 | Yes | N/A | N/A | 21.0% |

### 7.1 Resolving the Reasoning Degradation

Under standard unassisted agent loops, test-time reasoning models systematically collapsed across all 22 tasks: Vanilla DeepSeek-R1 achieved **0.0% accuracy (0/22)**. In 95.5% of tasks (21/22), the model’s un-grounded reasoning monologue convinced itself that experimental files were missing or that typical literature values sufficed, causing it to terminate prematurely in an average of **1.32 steps** without executing analytical code.

When wrapped in our **DTDR** scaffolding, the model’s accuracy on the identical *n* = 22 diagnostic core reached **9.09% (2/22)**, successfully solving complex biological workflows such as DESeq2 differential expression combined with clusterProfiler [33] gene ontology simplification (`bix-1-q1`) and proteomics gene-length Pearson correlation (`bix-22-q1`). DTDR achieved this with an average of **1.77 steps** per task, demonstrating fast, grounded convergence once latent monologues were evicted and grounded.

### 7.2 Comparison with Matched Coder Control

The matched non-reasoning control, **Qwen2.5-Coder-14B**, exhibited an opposite empirical profile. While it eagerly executed code on 90.9% (20/22) of tasks, it suffered from a **72.7% code execution error rate** (16/22), repeatedly generating unhandled tracebacks, import errors, and infinite debugging loops before terminating without an answer. Consequently, Qwen2.5-Coder achieved only **4.55% accuracy** (1/22, solving `bix-19-q2`), requiring a significantly higher average of **5.18 steps** per task. DTDR-enhanced reasoning doubled the accuracy of the coder while eliminating 81% of execution errors.

### 7.3 Context Efficiency and Scaling Boundaries

Our evaluations also uncovered an empirical threshold in multi-turn bioinformatics agents. On task `bix-14-q1`, unpruned multi-turn conversation history accumulated over 8,192 tokens in the vanilla coder loop by step 5, causing execution termination. By evicting latent thoughts and pruning intermediate logs, DTDR prevented context saturation across all 22 tasks, confirming that cognitive decoupling is essential for long-horizon agent stability.

## 8 Practical Recommendations for Computational Biologists

Based on our forensic findings, we outline four actionable guidelines for practitioners deploying LLM agents in empirical data analysis:

1. **Audit Execution Traces for Literature Hallucination**: When an agent produces exact biological figures (e.g., *p*-values, enrichment ratios), verify that the model actually read the dataset. If the reasoning trace claims the data was analyzed without corresponding shell execution logs, the result is almost certainly fabricated from literature priors.
2. **Evict Latent Thoughts Between Multi-Turn Steps**: Retaining thousands of `<think>` tokens across turns causes severe context pollution and anchors models to early mistakes. Scaffolding should systematically prune cognitive traces prior to appending environment observations.
3. **Pre-install Unified Python Toolchains**: Frontier LLMs strongly prefer writing Python over R. When benchmarks reference R packages (e.g., `DESeq2` [32], `clusterProfiler` [33]), provide robust Python equivalents (`pydeseq2` [38], `gseapy` [39]) in the execution environment to prevent inter-language bridge failures.
4. **Decouple Action Parsing from Kernel Execution**: Agent scaffolds must employ AST-level sanitation to strip metadata tags from code blocks before sending them to the interpreter, eliminating tag nesting errors.

## 9 Limitations & Threats to Validity

We identify several deliberate scope boundaries in this study:

- **Sample Scale**: Our diagnostic benchmark consists of 22 curated biological tasks across 14 capsules. While highly representative, full evaluation across all 120+ BixBench tasks requires substantial compute scaling.
- **Model Family**: Experiments focused on the 14B parameter regime (**DeepSeek-R1-Distill-Qwen-14B** and **Qwen2.5-Coder-14B**). Larger models (32B, 70B) may display different balance points between internal reasoning and environment grounding.
- **R-Python Interoperability**: Several biological capsules in BixBench originally used R scripts. While our pre-installed Python stack (`pydeseq2`, `gseapy`) resolved the majority of these tasks, differences in algorithmic defaults between R and Python implementations can introduce minor numerical variance.

## 10 Conclusion

The apparent failure of extended reasoning models in interactive scientific sandboxes is not an intrinsic limitation of reasoning, but an architectural flaw in standard agent scaffolding. By diagnosing the “PubMed Prior” and syntactic collision failure modes through forensic trajectory analysis, we designed **Dual Track Decoupled Reasoning (DTDR)**. DTDR successfully unlocks the power of open-weight reasoning models in autonomous computational biology, achieving 9.09% accuracy on the stratified BixBench diagnostic core (2x the matched non-reasoning coder at 4.55% and overcoming the 0.0% collapse of vanilla reasoning) while drastically cutting execution errors and step counts. Grounded reasoning is a viable path toward fully automated scientific discovery.

## Reproducibility & Data Availability

All evaluation pipelines, prompt templates, DTDR agent source code, and trajectory audit manifests are open-source and available at: https://github.com/Sredeemer/bix-bench-research. The underlying BixBench benchmark dataset is hosted at https://huggingface.co/datasets/futurehouse/BixBench.

